# Detecting ataxia using an automated analysis of motor coordination and balance of mice on the balance beam

**DOI:** 10.1101/2023.07.03.547476

**Authors:** Lucas Wahl, Fabian M.P. Kaiser, Mieke Bentvelzen, Joshua White, Martijn Schonewille, Aleksandra Badura

## Abstract

**Background:** The balance beam assay is a well-known paradigm to assess motor coordination in mouse models of neurodegenerative diseases. Classically, these experiments are quantified using manual annotation, which is time-consuming and prone to inter-experimenter variability.

**Methods:** We present an open-source analysis pipeline that allows for the automated quantification of motor function in mice crossing the balance beam.

**Results:** Using an established *Pcp2-Ppp3r1* ataxia mouse model, we validated the analysis pipeline by comparing the motor performance of *Pcp2-Ppp3r1* animals to their wildtype littermates. *Pcp2-Ppp3r1* animals showed a significant increase in the number of missteps and increased time to traverse the beam. Moreover, we compared the results of the automated classification of missteps and stops to that of 3 independent observers, which showed no significant differences between the classifier and the cumulative observer score for missteps.

**Conclusion:** We show that our pipeline can reliably report crossing time and missteps, offering a high-throughput, automated option for the analysis of balance beam data.

**Method summary:** Our method consists of an easy-to-follow and low-cost manual for building a balance beam setup and an analysis pipeline of mouse movement while crossing the beam. We present a MATLAB script combining FFMPEG and ImageJ (using the MIJ package) to pre-process the videos and extract the time for the mouse to cross the beam. Mice movements on the beam were then tracked using motr and stops and missteps were extracted using a trained JAABA classifier. The trained classifier, the code and all accompanying files were deposited in an open source repository

## Introduction

Mouse models of neurodegenerative disorders such as ataxia are indispensable for a better understanding of the mechanistic underpinnings of loss of motor coordination and balance. Until recently, the analysis of experimental data to quantitatively assess motor deficits has mostly relied on manual scoring by observers. Manual analysis of animal behaviour is time-consuming, arduous, labour-intensive, and subjective, often resulting in poor reproducibility across different observers due to inter-experimenter variability [1–5]. Machine learning and deep neural networks have greatly accelerated the analysis of animal behaviour across disciplines and provide standardised and objective results [6]. Such analysis strategies often rely on the training of automated classifiers by experienced investigators to annotate pre-defined behavioural or motor patterns until the software can reliably identify those without additional user input [3, 7–14].

The balance beam assay has emerged as a widely used and highly sensitive paradigm to study motor coordination and balance in mice [15]. The primary measurements of the assay are the time needed to traverse a beam that connects two platforms and the number of missteps the animals make during that period. The incentive to traverse the beam is created by either placing food or the animal’s homecage on the other platform. The task can be adapted by using flat or round beams of decreasing diameter or by varying the angle of the beam between the two platforms. The balance beam has proven to be more sensitive than the rotarod - another classic paradigm for motor coordination and balance assessment [16].

Other than the automated analysis of gait rhythmicity [17], the analysis of balance beam data has thus far been limited to the manual scoring of missteps. Here we report the development of an open-source analysis pipeline that we generated using the Janelia Automatic Animal Behavior Annotator (JAABA) [7], which reliably reports crossing time and missteps during the balance beam experiment. Using male and female adult *Pcp2-Ppp3r1* mice with Purkinje cell-specific knock-out of calcium/calmodulin-activated protein-phosphatase-2B (PP2B) on the *C57BL/6J* background - an established ataxia mouse model [18] - we provide proof-of-principle that the pipeline can discern phenotypic differences between ataxic mice and their wildtype littermates.

## Material and Methods

### Mice

Calcium/calmodulin-activated protein phosphatase 2B, also known as calcineurin, PP2B or protein phosphatase 3 (Ppp3r1), was selectively deleted from Purkinje cells with the Cre-loxP-system, using mice with loxP sites flanking the regulatory subunit,CNB1 (*Ppp3r1*^*tm1Stl*^) [19] and mice carrying the L7-Cre transgene (*Tg(Pcp2-Cre)3555Jdhu*) [20] as described in [18], to generate *Pcp2-cre*^+/-^-*Ppp3r1*^*fl/fl*^ mice (‘*Pcp2-Ppp3r1* mice’, n = 9) and *Pcp2-cre*^-/-^-*Ppp3r1*^*fl/fl*^ mice (hereafter referred to as wildtype animals, n = 8).

All mice were maintained on a C57BL/6J background. Additionally, 28 adult wildtype C57BL/6J animals were used for classifier training. Experimental animals were between 13 and 19 weeks of age (114 ± 17 days) and housed with food and water available *ad libitum* in a 12:12 hours light/dark cycle. All experiments were approved by local and national animal ethical committees (CCD approval: AVD1010020197846).

### Balance Beam setup

A balance beam setup was constructed using commercially available supplies. The setup was constructed with two opposing metal platforms positioned 50 cm above an acrylic glass plate with a distance of 86 cm between platforms (Figure 1A). Round aluminium beams of varying diameters (12/10 mm) were clamped below each platform with red markings on the beam indicating the start and end position of a section that measured 70 cm in length, thereby leaving 8 cm distance from the start and end positions on the track to each platform (Figure 1B). The 8 cm gap was excluded from analysis because mice generally paused to prepare for entering or exiting the beam on this section. A moveable aluminium arm with two joints that allows for adjustment of the camera was inserted at one side of the setup and the camera was placed at the same height as the beam. Recording was performed with a GoPro X HERO7 Black, capturing 29.97 frames per second at a resolution of 2704 × 1520 pixels in linear mode and MP4 format. To improve contrast, a white board was placed behind the setup (for white mice we recommend using a black board).

**Figure 1.**
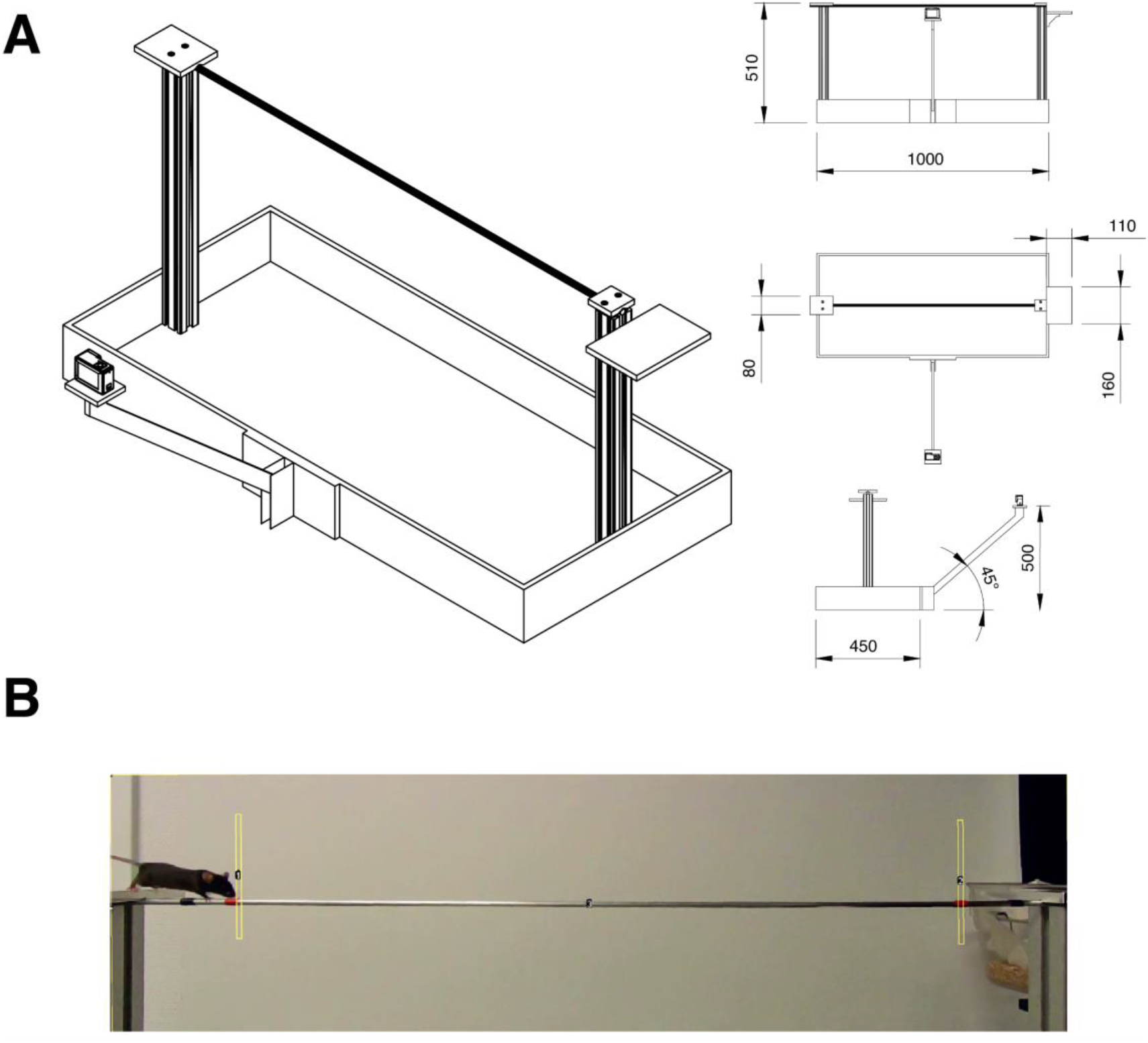
Setup standardisation. (**A**) Schematic of the balance beam apparatus used to quantify locomotor function in mice. Shown are the isometric view (left) and orthographic projection with size indicators. Measurements are shown in millimetres. (**B**) Side view of the beam in the balance beam test. Yellow rectangles indicate user-defined start and stop areas used in measuring behaviours and time to traverse the beam.

### Experimental design

Mice were habituated to the experimenters - who remained blinded to the genotype - by handling for 3-5 minutes per day for three consecutive days. Experiments were done in batches of 8 mice at the same time (8 mice per week), resulting in a total of 1 hour per batch handling per day. During the course of the experiment, mice were allowed to acclimate to the room for 1 hour prior to the beginning of the experiment. On day 1, mice were placed into the experimental cage with bedding of the home cage for 10 min prior to being placed on the platform on the opposite pole of the setup. On day 1 and day 2, mice were trained to traverse the balance beam on beams of decreasing diameter in three consecutive trials per beam. On day 1, beams with diameters of 12 mm and 10 mm were used. On day 2, mice were challenged to walk on beams with diameters of 10 mm, 8 mm, and 6 mm. *Pcp2-Ppp3r1* mice were unable to traverse the 8 mm and 6 mm beams, therefore only the 12 and 10 mm data is included in the results. On day 3, mice were evaluated on all beams in the order of decreasing diameter. After three consecutive trials on the same beam, mice were placed into the home cage and the setup was cleaned with 70% ethanol before the subsequent mouse was tested. Each mouse was given 1 min of rest after each trial. If mice failed three times during consecutive trials, the experiment was terminated and mice were tested on additional beams. At the beginning of each trial, a piece of paper displaying information about the animal, the beam diameter, and the current trial was briefly recorded. Afterwards, the animal was placed on the starting platform and allowed to traverse the beam. The arms of the investigator did not enter the field of view of the camera. Once the animal had reached the platform on the opposite side of the setup, recording was stopped. If the animal fell during the course of a trial, the trial was repeated.

### Pre-processing of videos

All subsequent analysis steps were performed on a computer equipped with an AMD Ryzen Threadripper 1950X 16-Core 3.4 GHz Processor with 16 cores and 64 GB RAM, using Windows 10 Pro 64-bit as the operating system. In the first step after data acquisition, videos were pre-processed in order to create files of the adequate format and length as well as to reduce the size, by integrating FFMPEG (https://ffmpeg.org/) and ImageJ [21] into MATLAB using the Image J MIJ package (https://imagej.net/plugins/miji). Using FFMPEG, videos were first cropped to cover the entire beam length and remove unnecessary areas above and below the beam. The video file was subsequently converted to the uncompressed AVI format to render it compatible with ImageJ. Within ImageJ, three regions of interest (ROI) were marked in the standardised video format to label the entire beam as well as the ROIs that designated the start and finish on the beam. The video was then trimmed based on the increase in signal in the start and finish ROIs as the mice traverse the beam in order to remove video segments during which mice were not on the beam, and a csv file with start and stop frames is created from which the time to traverse the beam can be calculated. After this step, the files were ready to be used as input for the tracking software to use for classifier training and behavioural quantification.

### Tracking

In the subsequent step, animals were tracked using the open-source software motr [2]. Motr tracks elliptical trajectories of mice in a closed environment with minimal user input. During the training, the user is asked to correct ellipses that are placed on the animal in order to compute the optimal segmentation threshold. The software uses previous frames to generate multiple hypotheses for the position of the mouse in the current frame and expectation maximisation algorithms are used to estimate the finite most probable position of the mouse in that frame.

### Manual labelling of videos

Observers received extensive instructions and training in scoring of behaviours on the balance beam from a skilled experimenter. Missteps were defined as “the foot coming off the beam”. This was accompanied with the downward motion of the hip.

### Training of the JAABA classifiers

In the final step, we used the open-source software JAABA [7] to train three independent classifiers to identify two common indicators of impaired motor coordination and balance: missteps and stops. Missteps or stops were defined as true positive events in the separate classifiers respectively and the sequences of the remaining video material were labelled as true negative events. JAABA uses supervised machine learning to create libraries with the designated motion patterns, processing information such as the position and angle of the animal as well as the trajectory data. We first implemented a batch-processing step in MATLAB for converting the trajectory data that was generated using motr into the correct input format for JAABA. We next trained a classifier to identify missteps, which we defined as slipping of the hindpaws that results in a downward motion of the hip, using 50 videos of adult male and female C57BL/6J and *Pcp2-Ppp3r1* mice, requiring ∼10 hours of training. In total, the classifier was trained on 3830 frames with 474 frames labelled as missteps and 3356 frames without such events. The minimum bout length was set to six frames (180 ms) by post-processing of the data.

Subsequently, we trained a classifier to recognize moments of stopping as the animals advanced on the beam. We used 80 videos of 28 adult male and female C57BL/6J mice to train the classifier. We defined stops as a pause in the movement of the animals over several sequential frames during the trial and trained the classifier with such as true positive events, using the remaining sequences - most including walking or missteps - as true negative events. In sum, the classifier was trained on 13235 frames with 6525 frames showing stops and 6711 frames without such events, requiring a total of ∼12 hours of training. We further set the amount of classifier training iterations to 200 and the cross-validation folds to 6. The minimum bout length was set to two frames (60 ms).

### Code accessibility

The code, classifiers and supporting files described in the article are freely available online at https://doi.gin.g-node.org/10.12751/g-node.el1sd7/.

## Results and Discussion

All experiments were performed on a standardised balance beam setup (Figure 1) built using readily available materials. In the Materials and Methods we provide details of setup assembly using commercially available materials from local hardware stores and a GoPro X HERO7 Black camera (Figure 1A). We estimate the construction time to be approximately 6 hours with estimated costs of ∼800 euro.

Next, we set up a standardised FFMPEG and FIJI analysis pipeline aimed at measuring the time required to traverse the beam as well as training a JAABA classifier to recognize two common indicators of impaired motor coordination and balance: missteps and stops (Figure 2). Specifically, in the first step after data acquisition, we processed videos using FFMPEG and ImageJ in order to acquire the time required for crossing the beam and to prepare the videos for the subsequent generation of trajectory data with motr [2]. To achieve this we wrote a custom MATLAB code using the MIJI plugin. We compared OptiMouse [22] and motr [2] - two open-source software packages for tracking the animals. In contrast to motr, tracking by OptiMouse was complicated by the frequent misidentification of the nose and tail, which caused the angle of the animal to turn regularly. This was most commonly observed when the tail of the animal escaped identification when positioned at the height of or below the beam. We therefore proceeded to use motr for the generation of the trajectory input for JAABA. Motr provides reliable, high-quality tracing data, but demands substantial processing power.

**Figure 2.**
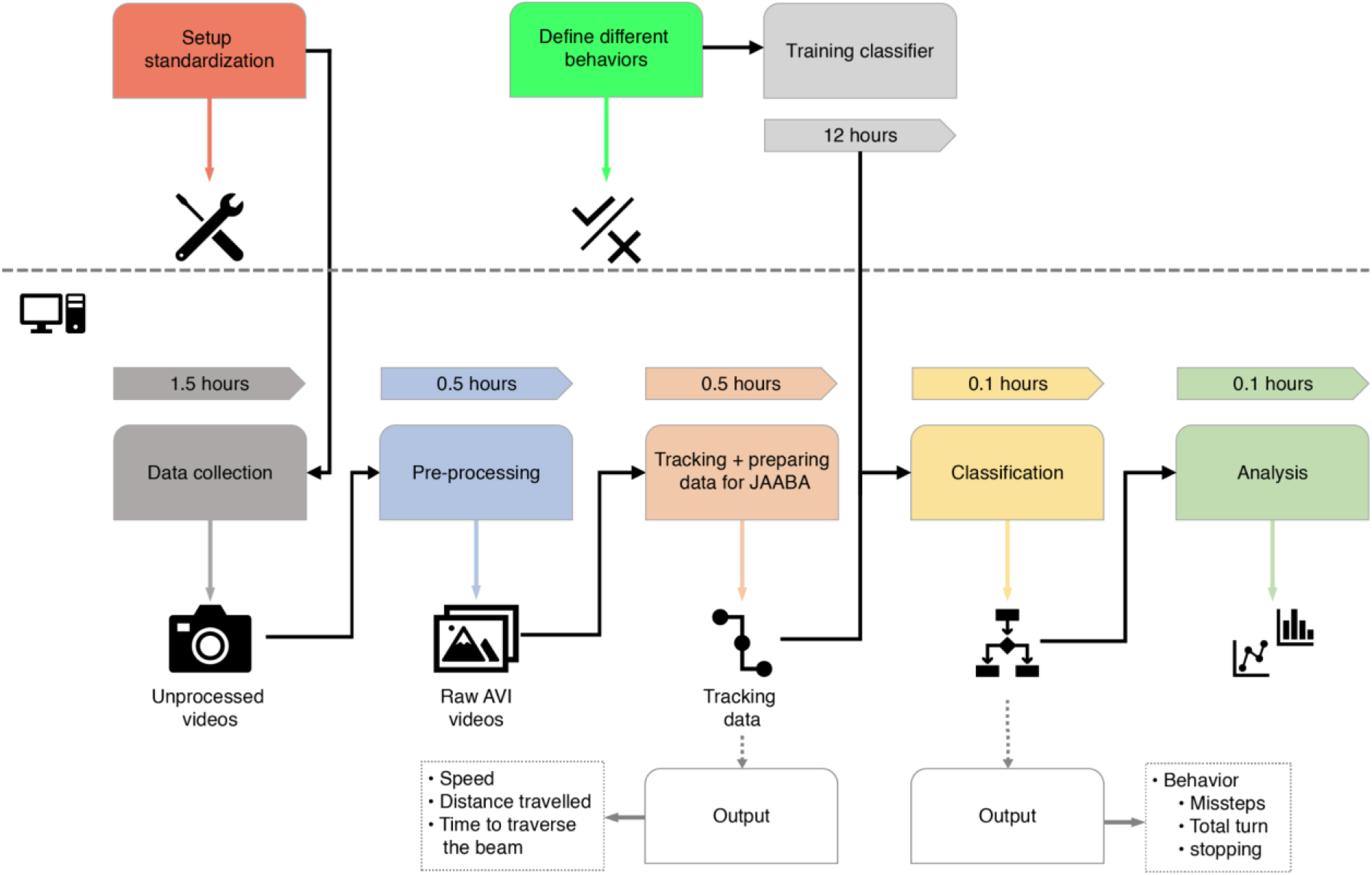
Pipeline for setting up standardised classification of the balance beam test for mice. The first step in setting up the balance beam test consists of building or acquiring the hardware apparatus, after which initial behavioural data can be collected. These videos have to be pre-processed in MATLAB to convert them for the open-source tracking software (Motr) and extract time of beam crossing using an ImageJ plug (MIJI). After the videos have been tracked, they can either be analysed with pre-existing classifiers, or used to train a new, custom classifier. Behaviours are then classified using the open-source software JAABA.

### Building an analysis pipeline to analyse missteps, stops and time to cross the balance beam

Using adult male and female C57BL/6J and *Pcp2-Ppp3r1* mice (a Purkinje cell-specific loss of the PP2B protein, necessary for long term potentiation - a known ataxic mouse model [18]) we trained two JAABA classifiers to recognize missteps (Figure 3A, left) and stops (Figure 3A, right), respectively, evaluating their accuracy using cross-validation. Cross-validation quantitatively assesses the accuracy of the classifier based on 1/7 folding by withholding a fraction of the dataset that is used for training of the classifier in order to validate its accuracy [7].

**Figure 3.**
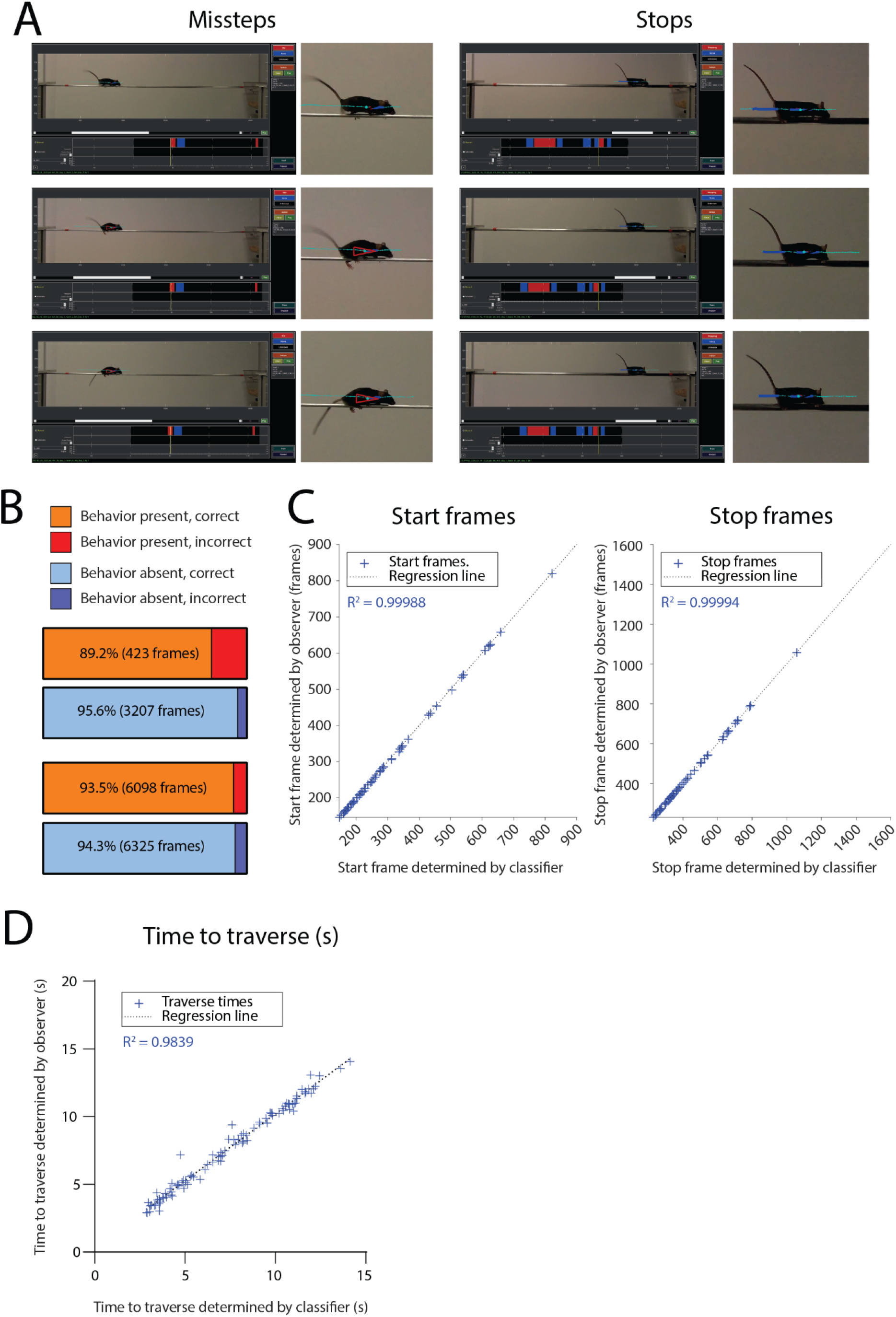
Classifier scoring and accuracy. (**A**) Sequence of selected frames of the classifier trained for the quantification of missteps in the balance beam test. Shown are the missteps (left) and stops (right). (**B**) Cross-validation results of manually annotated frames for the missteps (top), stops (bottom) classifiers. The orange bar represents correct annotation of frames containing the relevant behaviour, while red represents frames incorrectly scored as the relevant behaviour by the classifier. Light-blue bar represents correct annotation of non-behavior frames, whereas dark-blue shows incorrectly annotated non-behavior frames. (**C**) Manual scoring of start and stop frames in a set of 60 videos revealed highly correlated time to traverse the beam with ImageJ, as indicated by Pearson’s R square value. (**D**) Manual scoring of the time to traverse the beam in a set of 60 videos revealed highly correlated time to traverse the beam with ImageJ, as indicated by Pearson’s R square value.

We defined missteps as slipping of the hindpaws off the top of the beam, which was observed to be accompanied with a downward motion of the hip (Figure 3A, left). The corresponding classifier reached a sensitivity of 89.2% to identify such and a specificity of 95.6% to identify segments without missteps with a minimum length of six frames (180 ms) required for the identification (Figure 3B, top). Stops were defined as pausing of the animal for at least 2 frames (60 ms) before continuing forward locomotion. With regard to stops, the classifier reached a sensitivity of 93.5% to accurately identify those and a specificity of 94.3% to identify segments without stops (Figure 3B, bottom).

In order to determine whether the automated analysis pipeline was able to accurately determine the time needed to traverse the beam, we manually analysed 60 videos of adult male and female C57BL/6J mice. We observed a high correlation between the manual and automated analysis in determining the start (Figure 3C, left; R^2^ = 0.99988) and stop frame (Figure 3C, right; R^2^ = 0.99994), respectively, as well as the time to traverse the beam (Figure 3D; R^2^ = 0.8089). As mentioned above the time to traverse the beam was determined during the pre-processing step using the ImageJ MIJI plugin and a custom MATLAB code.

### Comparison of classifiers to observers’ scores

We subsequently compared the classifier performance to three observers that were blinded to the genotype of the animals and introduced the underlying principle of the assay. We used 102 videos of male and female *Pcp2-Ppp3r1* adult mice with Purkinje cell (PC)-specific deletion of the PP2B protein, resulting in a lack of parallel fibre-to-PC long-term potentiation [18, 23] and their wildtype littermates for this task. With regard to missteps, we observed substantial variation in scoring by manual observers (Figure 4A). Specifically, whereas observer 1 and 3 showed consistent results across genotypes, observer 2 experienced great difficulty in identifying missteps, even after receiving extensive instructions and training in scoring from a skilled experimenter. This finding underlines the known problem of inter-experimenter variability in the manual analysis of behavioural and motor coordination experiments with rodents.

**Figure 4.**
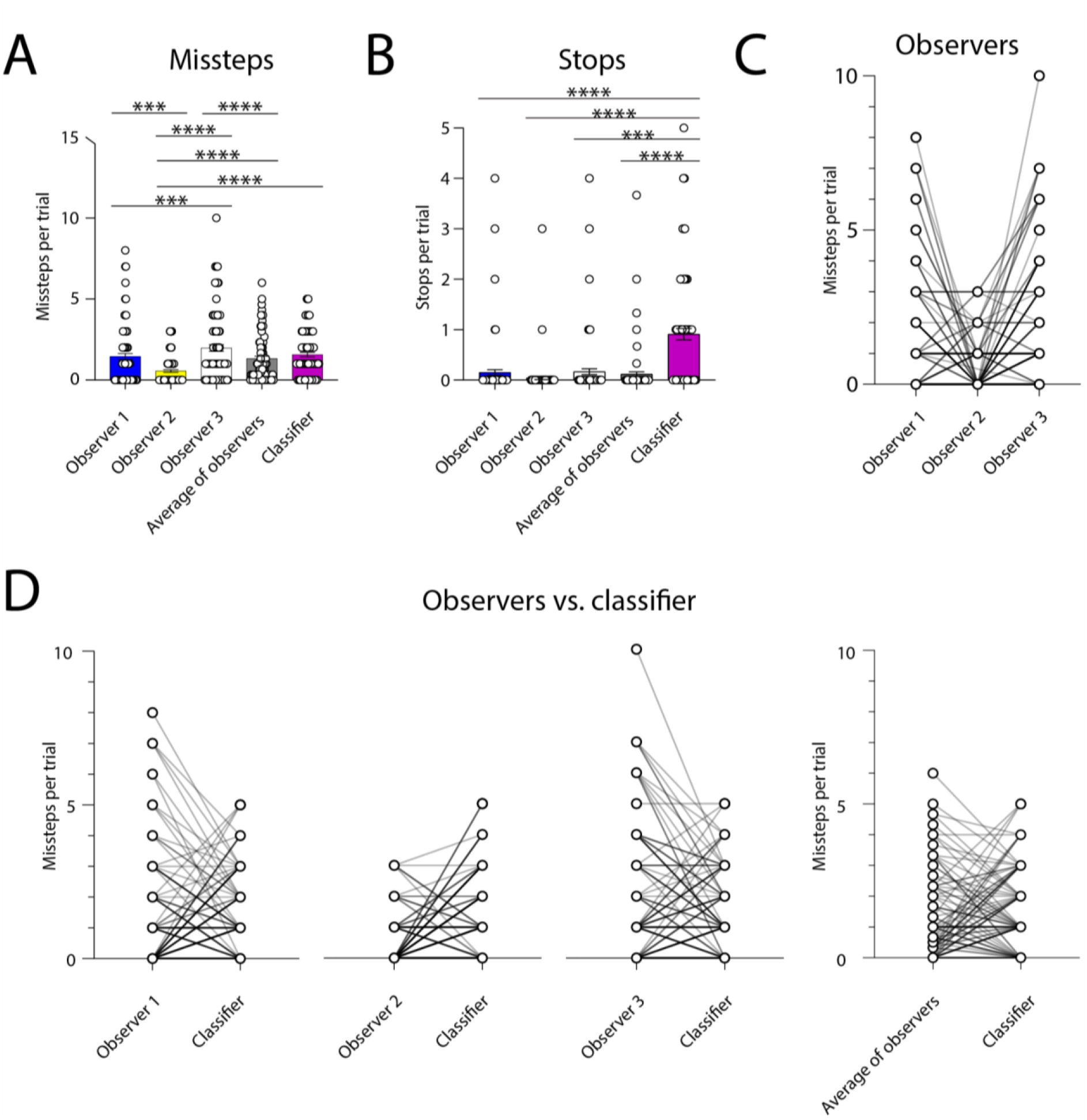
Classifier validation. (**A**) Number of missteps in 102 videos, manually scored by three observers as well as a trained classifier. (**B**) Number of stops in 102 videos, manually scored by three observers as well as a trained classifier. (**C**) Ranking correlation between scoring of the number of missteps per trial by observers. (**D**) Ranking correlation between scoring of the number of missteps per trial by observers and the classifier. Data presented as mean with SEM. Missteps and stops, Kruskal-Wallis test. *p ≤ 0.05, **p ≤ 0.01, ***p ≤ 0.001, n = 17 mice.

When comparing the identification of stops, the classifier identified more stop events than the three observers (Figure 4B). We have set the minimum number of frames per stop to two, thereby also defining short moments of pausing as stops. These short bouts may have been difficult to discern for the observers. Nevertheless, mice that have motor coordination problems often stop for short periods of time, which might elude the observers. We therefore opted to keep this increased sensitivity of the classifier to capture the whole range of motor deficits of the *Pcp2-Ppp3r1* mice. Observers showed large variability in the ranking of mice when scoring missteps per trial (Figure 4C), with overall consistent results with the classifier (Figure 4D,E).

### Automated classification of behaviours and speed using an ataxia mouse model

In order to determine whether the classifiers were capable of reliably identifying differences in motor coordination and balance - as reflected by quantification of missteps and stops - between ataxic and wildtype mice, we evaluated the performance of the *Pcp2-Ppp3r1* mice and wildtype littermates, with the use of automated classification. Given the severe motor coordination impairments of the *Pcp2-Ppp3r1* animals, we were only able to evaluate their performance on 12 mm and 10 mm beams. *Pcp2-Ppp3r1* mice showed a significant increase in missteps per trial on both the 12 mm beam (Figure 5A, left; p = 0.007) and a trend for the same pattern on the 10 mm beam (Figure 5A, right; p = 0.11), confirming the ataxic phenotype of *Pcp2-Ppp3r1* mice. No significant difference was found when analysing the number of stops per trial between wildtype and *Pcp2-Ppp3r1* animals (Figure 5B; 12 mm beam, p = 0.3000; 10 mm beam, p = 0.2240). In order to determine whether the animals exhibit altered locomotor patterns on the balance beam, we used our automated FFMPEG and FIJI analysis pipeline to measure the speed at which animals traverse the beam. *Pcp2-Ppp3r1* animals took significantly longer to traverse both the 12 mm beam (Figure 5C, left; p < 0.0002) and the 10 mm beam (Figure 5D, right; p < 0.0001), unveiling another metric that differs between wild-type and *Pcp2-Ppp3r1* animals. The number of missteps scored by the classifier and the time needed to traverse the beam were highly correlated for the 12 mm beam (Figure 5D, left) as well as the 10 mm beam (Figure 5D, right), showing that the reduced speeds are due to the ataxia phenotype and not a coping mechanism to reduce errors on the beam.

**Figure 5.**
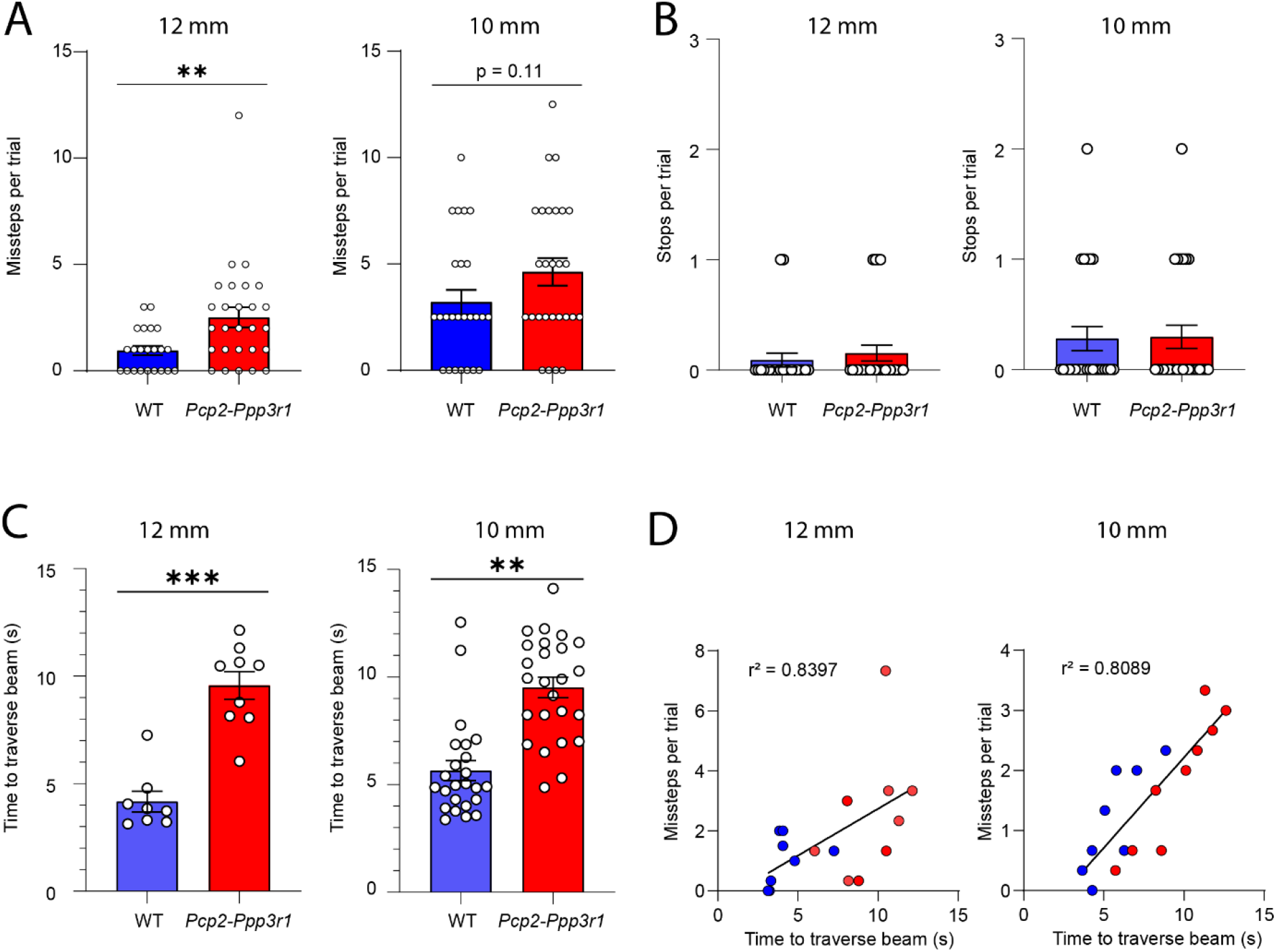
Automated behavioural classification of missteps, stops and crossing time in the balance beam test. Automated classification of behaviours is shown for *Pcp2-Ppp3r1* mice (red) and wildtype littermates (WT, blue) during the balance beam test). (**A**) Number of missteps per trial. (**B**) Number of stops per trial. (**C**) Time to traverse the beam per trial. (**D**) Correlation between the missteps per trial and the time needed to traverse the beam. Data presented as mean with SEM, number of missteps (12 mm and 10 mm), two-tailed t test; number of stops (12 mm), Mann-Whitney test; number of stops (10 mm), two-tailed, unpaired t test; time to traverse the beam (12 mm), Mann-Whitney test; time to traverse the beam (10 mm), two-tailed t test. *p ≤ 0.05, **p ≤ 0.01, ***p ≤ 0.001, n = 8 for WT mice and n = 9 for *Pcp2-Ppp3r1* mice.

## Conclusion

The analysis of rodent behaviour and motor coordination experiments has been facilitated and accelerated by the introduction of machine learning algorithms and deep neural networks, generating quantitative, reliable, and high-throughput measurements without inter-experimenter variability. Here we describe and validate an automated analysis pipeline for the balance beam assay - a well-established and sensitive paradigm for the analysis of motor coordination and balance deficits in rodents. Our pipeline reports the time needed to cross the beam as well as the amount of missteps and stops. This pipeline accelerated the analysis and reliably identified motor coordination deficits of the *Pcp2-Ppp3r1* animals on the balance beam. It offers a cost-effective and high-throughput alternative to manual scoring of behaviour. Finally, our analysis relies on open-source software, and can be easily applied in other experimental setups.

## Executive summary

### Background

We developed a cost-effective and high-throughput alternative to manual scoring of motor function on the balance beam.

### Methods

We validated the open-source analysis pipeline by comparing its output to three independent observers and quantifying the differences in motor function between wildtype and *Pcp2-Ppp3r1* animals (an established ataxia model), on the balance beam.

### Results

The pipeline could reliably report crossing time and missteps. Further using our pipeline, we reproduced previously published results showing that the *Pcp2-Ppp3r1* animals exhibit a significant increase in the number of missteps and increased time needed to traverse the beam.

### Conclusion

Our setup and analysis pipeline offers a cost-effective and high-throughput option with increased inter-experimenter and intra-experimenter reliability for the analysis of balance beam data.

## Acknowledgements

The authors are grateful to Maaike Imthorn for assistance with animal experiments.

